# Modelling the population dynamics of *Plasmodium falciparum* gametocytes in humans during malaria infection

**DOI:** 10.1101/641472

**Authors:** Pengxing Cao, Katharine A. Collins, Sophie Zaloumis, Thanaporn Wattanakul, Joel Tarning, Julie A. Simpson, James S. McCarthy, James M. McCaw

**Author notes:** Address correspondence to Pengxing Cao or James M. McCaw.

## Abstract

Every year over two hundred million people are infected with the malaria parasite. Renewed efforts to eliminate malaria has highlighted the potential to interrupt transmission from humans to mosquitoes which is mediated through the gametocytes. Reliable prediction of transmission requires an improved understanding of *in vivo* kinetics of gametocytes. Here we study the population dynamics of *Plasmodium falciparum* gametocytes in human hosts by establishing a framework which incorporates improved measurements of parasitaemia in humans, a novel mathematical model of gametocyte dynamics, and model validation using a Bayesian hierarchical inference method. We found that the novel mathematical model provides an excellent fit to the available clinical data from 17 volunteers infected with *P. falciparum*, and reliably predicts observed gametocyte levels. We estimated the *P. falciparum*’s sexual commitment rate and gametocyte sequestration time in humans to be 0.54% (95% credible interval: 0.30-1.00) per life cycle and 8.39 (6.54-10.59) days respectively. Furthermore, we used the data-calibrated model to predict the effects of those gametocyte dynamics parameters on human-to-mosquito transmissibility, providing a method to link within-human host kinetics of malaria infection to epidemiological-scale infection and transmission patterns.

---

Malaria is a mosquito-borne parasitic disease caused by protozoan parasites of the *Plasmodium* genus. It is estimated to have caused approximately 219 million new cases and 435,000 deaths (primarily due to *Plasmodium falciparum*) in 2017^1^. New tools will be required to achieve the ambitious goal of malaria elimination. Among tools proposed are novel interventions to block transmission of infection from human hosts to vector mosquitoes^2^. Falciparum malaria is transmitted from the human host to the mosquito when terminally differentiated male and female sexual stages of the parasite, called gametocytes, are taken up by the female *Anopheles* mosquito during a blood meal^3,4^. The level of gametocytes in the blood, often referred to as gametocytaemia, is highly associated with the probability of human-to-mosquito transmission^5,6^. Gametocyte levels below a certain threshold (i.e. < 1000 per mL blood) minimise the probability for a mosquito to take up both a male and female gametocyte during a blood meal, necessary to propagate infection^7^. An accurate understanding of the kinetics of gametocytes in human hosts is essential to predict the probability of transmission. A mathematical model that accurately captures the processes that give rise to observed gametocyte kinetics would be an important predictive tool to facilitate the design of effective intervention strategies.

Nonetheless, there is significant uncertainty surrounding fundamental aspects of *Plasmodium falciparum* gametocyte population dynamics in humans. Parameters such as how many gametocytes are produced during each asexual life cycle, the period of time in which early gametocytes disappear from the circulation before mature gametocytes appear (referred to as sequestration), and the period in which gametocytes circulate are poorly quantified. These gaps in understanding are due to a range of technical and logistic limitations. The first is the relatively poor sensitivity of the standard diagnostic test, namely microscopic examination of blood films. *In vivo* estimates of gametocyte kinetic parameters have been primarily based on historical data from neurosyphilis patients whose illness was treated with so-called malariotherapy^8,9^. In these studies, the limit of quantification was approximately 10^4^ parasites/mL blood, at least two orders of magnitude higher than that of current quantitative PCR (qPCR) assays^10^. This high limit of quantification prevents an accurate estimation of the onset of the emergence of both asexual parasites and mature gametocytes in peripheral blood. The second limitation is that the available estimates of gametocyte population dynamics parameters based on *in vitro* cultures^11,12^ may not applicable to natural infection with *P. falciparum* gametocytes due to the *in vitro* conditions that may not mimic what occurs in the human host^3^.

Recent advances in experimental medicine using volunteer infection studies (VIS), otherwise known as controlled human malaria infection (CHMI) studies, allow prospective study design and data collection with the explicit aim of collecting *in vivo* data^13^, in particular an improved quantification of *P. falciparum* gametocyte kinetics by qPCR applied in a novel VIS^7^. Furthermore, the models and fitting methods used in the neurosyphilis patient studies have been superseded for parameter estimation by increasingly sophisticated within-host models^14^ and improvements in computational algorithms for Bayesian statistical inference^15^. Therefore, there is an emerging opportunity to improve our quantitative understanding of the population dynamics of *P. falciparum* gametocytes in human hosts by combining the novel VIS data and advanced modelling approaches.

In this paper, we developed a novel mathematical model of gametocyte dynamics, fitted the model to the VIS data and estimated the gametocyte dynamics parameters using a Bayesian hierarchical inference method (details are given in the Materials and Methods). We demonstrate that the data-calibrated model can reliably predict the time course of gametocytaemia and thus should form an essential part of modelling studies of malaria transmission.

## Results

### Model fitting and validation

The outcome variable used in model fitting was the total parasitaemia (total circulating asexual parasites and gametocytes per mL blood) collected from a previously published VIS^7^, with other measurements from the same study, such as the asexual parasitaemia (circulating asexual parasites per mL blood) and gametocytaemia (circulating female and male gametocytes per mL blood), used to validate model predictions.

The results of fitting the mathematical model to total parasitaemia data for all 17 volunteers are given in Figure 1 (upper panels) where 12 volunteers experienced recrudescence while the rest did not. The median of posterior predictions and 95% prediction interval (PI) were computed from 5000 model simulations based on 5000 samples from the posterior parameter distribution (see Materials and Methods). The results show that the model describes the total parasitaemia data very well (median of posterior predictions vs. observed data). Furthermore, the narrow 95% PI indicates a low level of uncertainty in predicted total parasitaemia.

**Figure 1:**
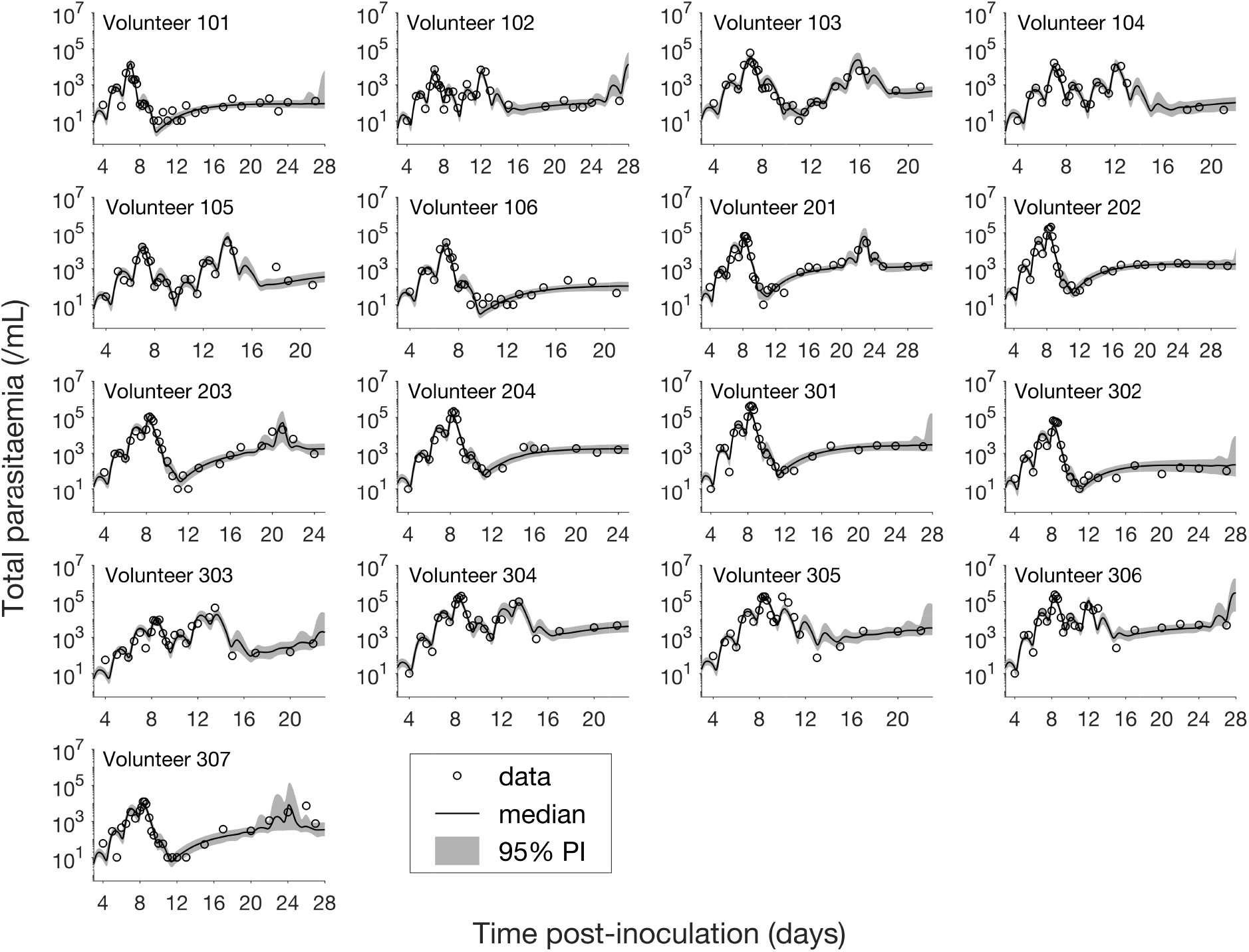
Results of model fitting for all 17 volunteers. Data are presented by circles. The median of posterior predictions (solid line) and 95% prediction interval (PI, shaded area) are generated by 5000 model simulations based on 5000 samples from the posterior parameter distribution (see the Materials and Methods).

Having calibrated the model against total parasitaemia, the 5000 posterior parameter sets were used to calculate the median of posterior predictions and 95% PI of the asexual parasitaemia and gametocytaemia versus time profiles. Model predictions of the asexual parasitaemia and gametocytaemia for all 17 volunteers are shown in Figure 2 and Figure 3 respectively (curves: median prediction; shaded areas: 95% PI) and are compared to the asexual parasitaemia and gametocytaemia data (circles). We emphasise that this was not a fitting exercise, rather an independent validation of the calibrated model.

**Figure 2:**
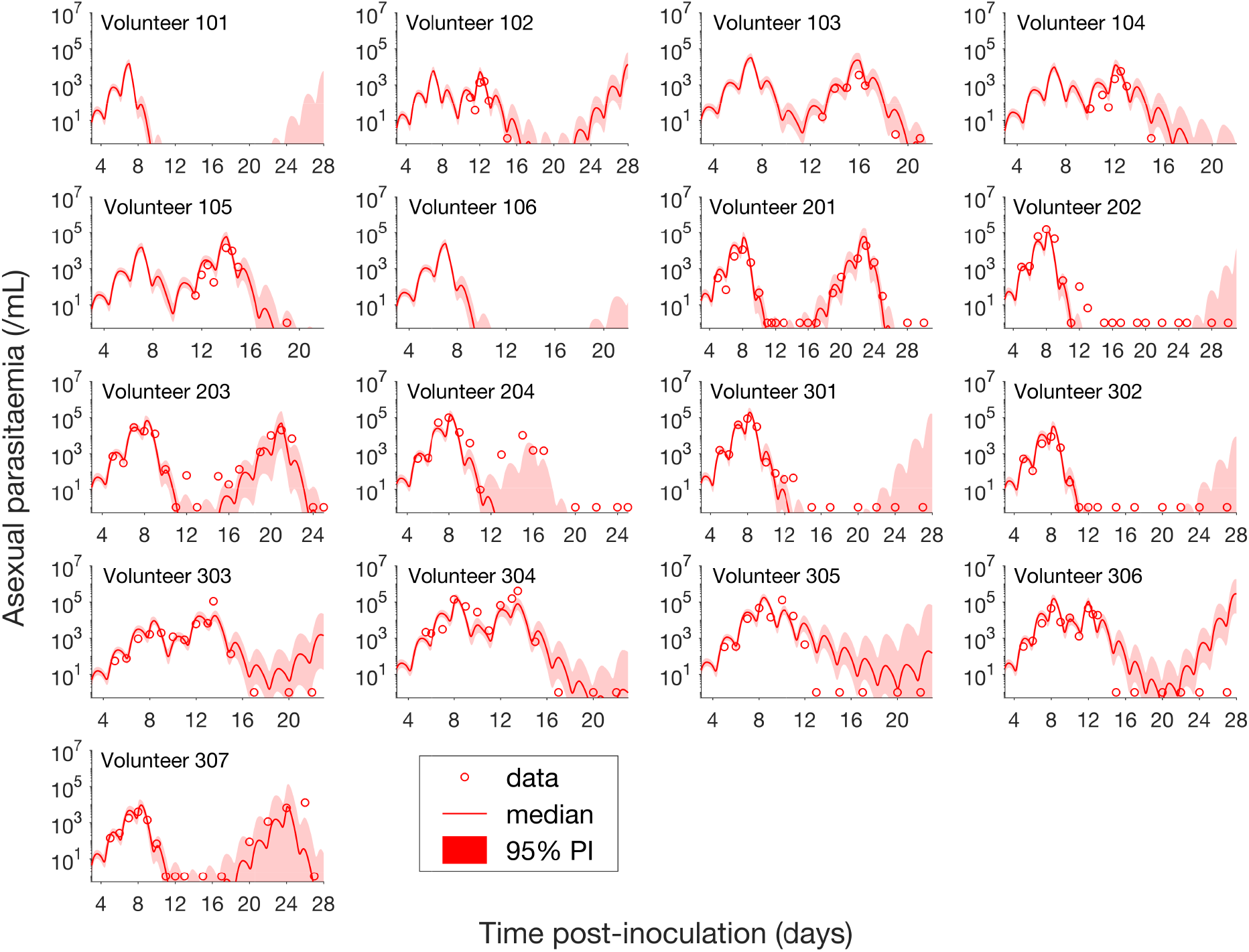
Comparison of model predictions and clinical data for the asexual parasitaemia for all 17 volunteers. Data are presented by circles. The median of posterior predictions (solid curve) and 95% PI (shaded area) are generated by 5000 model simulations based on 5000 samples from the posterior parameter distribution (see the Materials and Methods). Note that one asexual parasite/mL was the level of detection and no data are available for Volunteer 101 and 106 to validate the model predictions.

**Figure 3:**
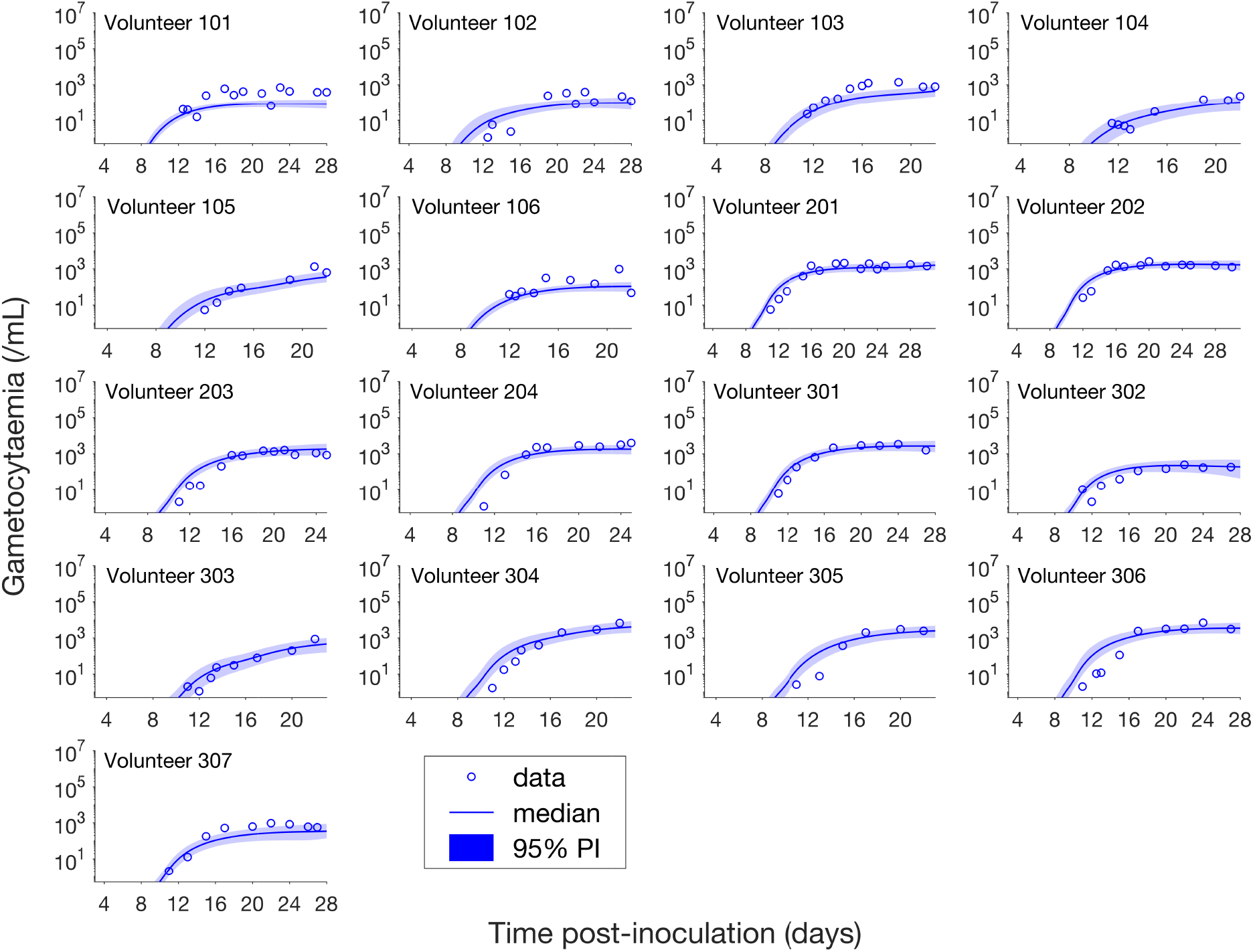
Comparison of model predictions and clinical data for the gametocytaemia for all 17 volunteers. Data are presented by circles. The median of posterior predictions (solid curve) and 95% PI (shaded area) are generated by 5000 model simulations based on 5000 samples from the posterior parameter distribution (see the Materials and Methods). Note that one gametocyte/mL was the level of detection.

For the majority of asexual parasitaemia data there is an excellent predictive capability for the model (see Figure 2), in particular the accurate predictions for both the recrudescent case where a portion of asexual parasitaemia data diverge from the total parasitaemia measurement (e.g. Volunteer 103, 104, 105, 201, 203, 304 and 307) and the non-recrudescent case where the posterior-median prediction curve (solid red curve) lies below the limit of detection (one asexual parasite/mL) (e.g. Volunteer 202, 301 and 302). However, there are some discrepant observations. The model under-predicts (see Volunteer 204) or over-predicts (see Volunteer 303, 305 and 306) a portion of the asexual parasitaemia data. Furthermore, for some volunteers such as 202, 301 and 302, the 95% PI (red shaded area) extends into the detectable region again after the asexual parasitaemia reaches below the detection limit, indicating that there was a small chance that the patients may have suffered a recrudescence during the observation period (of course, they did not) or after the observation period (although this predication cannot be evaluated because artemisinin combination therapy was given immediately after the period). Despite the above discrepant observations for asexual parasitaemia, the model predictions of gametocytaemia are very consistent with the data for all 17 volunteers, as shown in Figure 3.

### Estimation of gametocyte population dynamics parameters

The model calibration process provided the joint posterior density for the model parameters, which were used to estimate some key biological parameters governing the population dynamics of *P. falciparum* gametocytes (see the Material and Methods for details). As shown in Table 1, the sexual commitment rate – the percentage of asexual parasites entering sexual development during each asexual life cycle – is estimated to be 0.54% (95% credible interval (CI): 0.30%, 1.00%). This *in vivo* estimate is much lower than 11% that was estimated from *in vitro* data^11^. The gametocyte sequestration time is the average time that immature gametocytes (stages I—IV) cannot be observed in the peripheral circulation and was estimated to be 8.39 days (95% CI: 6.54, 10.59 days). The estimate for the circulating gametocyte lifespan is 63.5 days, with a broad 95% CI (12.7 to 1513.9 days) resulting from the long-tailed posterior distribution (see Figure S1, Supplementary Information) and is much longer than the previous *in vitro* estimate of 16—32 days^12^ (note that our lower bound of the 95% CI is lower than the *in vitro* estimated range). The wide estimate for the circulating gametocyte lifespan, and in particular the high upper bound of the 95% CI, is primarily due to the limited observation time in the VIS as explored in more detail in the Discussion.

**Table 1:** Estimates of some key biological parameters and comparison with the literature. The estimates of the biological parameters (middle column) are shown as the median and 95% credible interval (CI) of the marginal posterior parameter distribution (see Figure S1 in the Supplementary Information). Estimates from the literature (third column) are shown in the format of either “mean estimate (95% confidence interval)” or “mean estimate [minimum – maximum estimate]” or simply “a low estimate *–* a high estimate”. Note some quoted estimates are from studies of *P. falciparum* (*Pf*) from either *in vivo* or *in vitro* studies.

| Biological parameters (unit) | Median estimate (95%CI) | Estimates in the literature |
| --- | --- | --- |
| Sexual commitment rate (%/life cycle) | 0.54 (0.30, 1.00) | 11 (6.2, 15.8)<sup>11</sup> (*Pf*; *in vitro*)<br>0.64 [0.03 – 13.51]<sup>9</sup> (*Pf*; *in vivo*) |
| Gametocyte sequestration time (days) | 8.39 (6.54, 10.59) | 7.4 [4 – 12]<sup>9</sup> (*Pf*; *in vivo*) |
| Circulating gametocyte lifespan (days) | 63.5 (12.7, 1513.9) | 16 – 32<sup>12</sup> (*Pf*; *in vitro*)<br>6.4 [1.3 – 22.2]<sup>9</sup> (*Pf*; *in vivo*) |
| Parasite multiplication factor (/life cycle) | 21.8 (17.6, 26.9) | 10 – 33<sup>16</sup> (*Pf*; *in vivo*)<br>16.4 (15.1, 17.8)<sup>17</sup> (*Pf*; *in vivo*) |

As shown in Table 1, there are both similarities and differences in parameter estimates for *P. falciparum* based on our analysis and the analysis of historical neurosyphilis patient data^9^. We found that they exhibited very similar *in vivo* sexual commitment rate (median 0.54% vs. mean 0.64% with overlapping plausible ranges) and gametocyte sequestration time (median 8.39 days vs. mean 7.4 days with overlapping plausible ranges). On the other hand, we found that the circulating gametocyte lifespan for *P. falciparum* was much longer than that estimated from the neurosyphilis patient data^9^ (median 63.5 days vs. 6.4 days with very different ranges).

Finally, we provided an estimate for the parasite multiplication factor, an important parameter that quantifies the *net* replication rate of asexual parasites and also influences the rate of gametocyte generation. Our posterior-median estimate is 21.8 parasites per life cycle (95% CI: 17.6, 26.9), consistent with previous estimates which lie in the range 10—33^16^, and a bit larger than an updated estimate calculated from a pooled analysis of parasite counts from 177 volunteers infected with the same *P. falciparum* strain (16.4 parasites per life cycle)^17^.

### Predicting the impact of gametocyte kinetics on human-to-mosquito transmissibility

Having validated our mathematical model of asexual parasitaemia and gametocyte dynamics, we were able to predict how the gametocyte dynamics parameters influence the transmissibility of *P. falciparum* malaria from humans to mosquitoes in various epidemiological scenarios. In particular, we focused on the early phase of infection where the innate immune response is minimal and treatment has not been administered (in order to avoid complications that our mathematical model was not designed to capture). Two scenarios were considered:

- Predicting the potential infectiousness of newly hospitalised malaria patients for various values of sexual commitment rate and gametocyte sequestration time. In the model, gametocytaemia was assumed to be a surrogate of the potential infectiousness and newly hospitalised patients were assumed to exhibit a total parasitaemia of approximately 10^8^ parasites/mL based on published clinical data from Cambodia and Thailand^18^. As illustrated in Figure 4A, we simulated the model (for different sexual commitment rates and gametocyte sequestration times) and looked at the critical gametocytaemia level (indicated by *G*_*c*_) corresponding to the time when the total parasitaemia (wave-like black curves) first reached 10^8^ parasites/mL (their associations are indicated by the dotted lines and arrows).
- Predicting the non-infectious period of malaria patients for various values of sexual commitment rate and gametocyte sequestration time. In the model, the non-infectious period was defined to be time from the inoculation of infected red blood cells to the time when the gametocytaemia reach 10^3^ parasites/mL, which is a threshold allowing for human-to-mosquito transmission^7^. Note that this non-infectious period does not include the latent period due to the liver stage which should be considered if the starting time were taken as the inoculation by mosquito. As illustrated in Figure 4B, we simulated the model (for different sexual commitment rates and gametocyte sequestration times) and identified the critical time (indicated by *t*_*c*_) when the gametocytaemia (blue curve) first reached 10^3^parasites/mL (their associations are indicated by the dotted lines and arrows).

**Figure 4:**
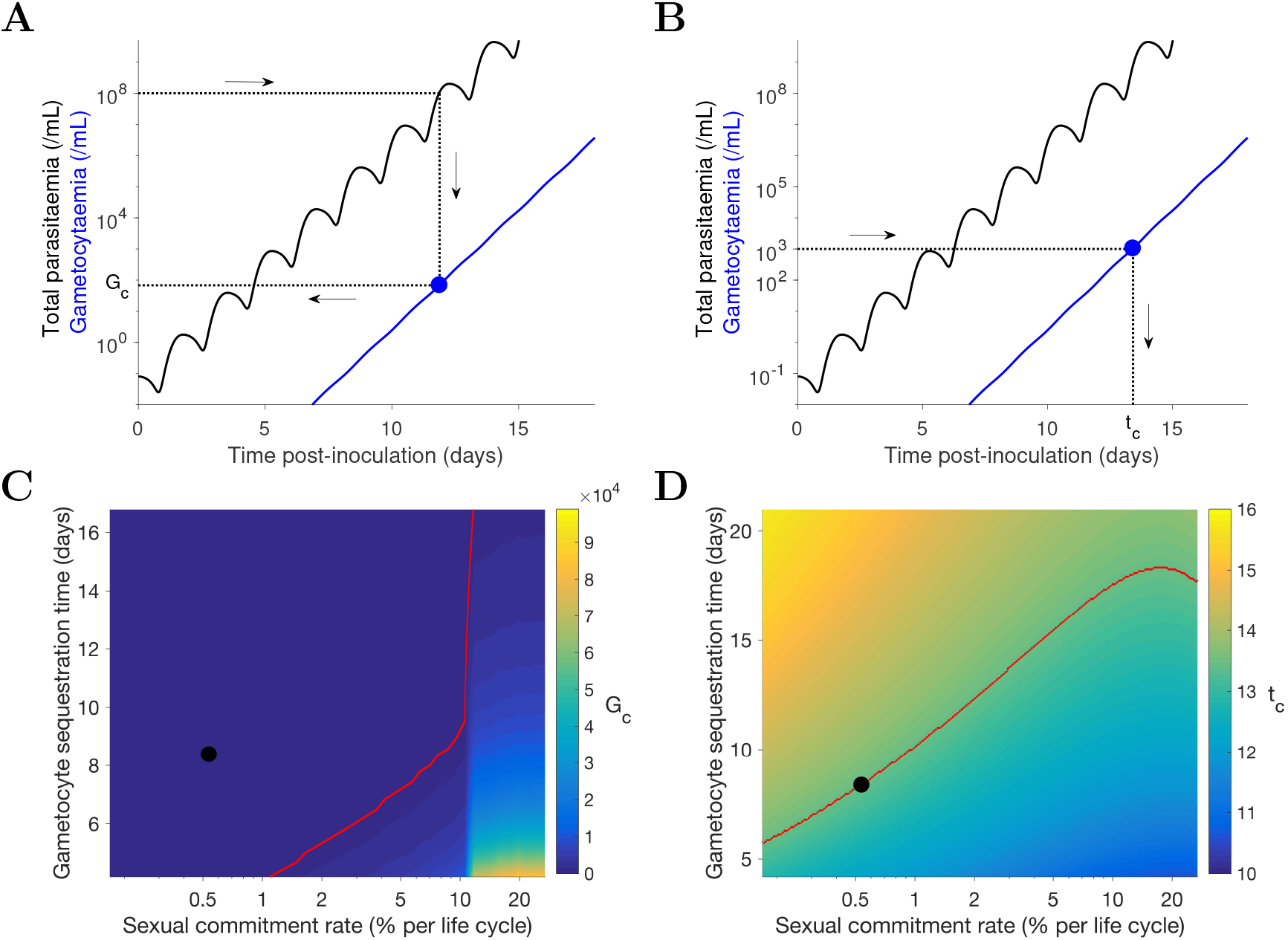
Simulation of two scenarios predicting the dependence of human-to-mosquito transmissibility on the sexual commitment rate and gametocyte sequestration time. (A) illustration of the first scenario: predicting the critical gametocytaemia level (indicated by *G*_*c*_) at the time when the total parasitaemia reach 10^8^ parasites/mL. (B) illustration of the second scenario: predicting the non-infectious period (indicated by *t*_*c*_), which is defined to be time from inoculation of infected red blood cells to the time when the gametocytaemia reach 10^3^ parasites/mL (a threshold for human-to-mosquito transmission^7^). (C and D) Heatmaps showing the dependence of the critical gametocytaemia *G*_*c*_ and the non-infectious period *t*_*c*_ on the sexual commitment rate and gametocyte sequestration time. The black dots represent the benchmark value obtained by simulating the gametocyte population dynamics model using the median estimates of the posterior samples of the population mean parameters (see the Materials and Methods for details). The red curves indicate the cases corresponding to gametocytaemia of 10^3^ parasites/mL.

Figure 4C shows that a higher sexual commitment rate or a lower gametocyte sequestration time leads to a higher gametocytaemia (*G*_*c*_) at the time of hospitalisation. The red curve in Figure 3C indicates the level curve of 10^3^ gametocytes/mL (i.e. the threshold for infectiousness as mentioned above) dividing the heatmap into two regions. To the left, *G*_*c*_ is below 10^3^ gametocytes/mL, suggesting clinical presentation precedes infectiousness, while to the right *G*_*c*_ is above 10^3^ gametocytes/mL and the converse applies. The *G*_*c*_ value obtained by model simulation using the median estimates of the population mean parameters (indicated by the black dot) is below 10^3^ gametocytes/mL, suggesting that newly hospitalized malaria patients are less likely to be infectious, and thus efforts to identify and treat infections in a timely manner may have a substantial impact in terms of reduced transmission potential.

Figure 4D reinforces the result in Figure 4C using the non-infectious period (*t*_*c*_). As the sexual commitment rate increases or the gametocyte sequestration time decreases, *t*_*c*_ decreases. However, for relatively large values of the sexual commitment rate (e.g. > 20%), we observed an increase in *t*_*c*_ as the sexual commitment rate increases (see the top-right corner of Figure 4D). This is because an increased fraction of sexual conversion can lead to both a decrease in the rate of asexual parasite growth and an increase in the number of sexual parasites, and the impact of the former more than counterbalances that of the latter.

## Discussion

We have developed a novel mathematical model of gametocyte dynamics, that integrates both multi-state asexual parasite’s life cycle and the development of gametocytes for the first time. Model parameters were estimated by fitting the model to data from 17 malaria-naïve volunteers inoculated with *P. falciparum*-infected red blood cells (3D7 strain). Compared to previous studies, our work is distinguished by three novel contributions: (1) the use of a prospectively planned clinical trial to collect more accurate quantitative data of parasite levels measured by qPCR; (2) the development of a novel population dynamics mathematical model which allows for robust and biologically-informed extrapolation and hypothesis testing/scenario analysis; and (3) the use of a Bayesian hierarchical inference method for model calibration and parameter estimation.

For gametocyte kinetic parameters, we found that our *in vivo* estimate of the *P. falciparum* sexual commitment rate was similar to that found in the neurosyphilis patient data^9^ but was much smaller than previous *in vitro* estimates (see Table 1). Novel VIS data using biomarkers specific to early sexual parasites (e.g. PfGEXP5^19^) may help to validate and improve our *in vivo* estimate. Furthermore, our *in vivo* estimate of the circulating gametocyte lifespan is substantially larger than previous *in vitro* estimates and is also larger than that found in the neurosyphilis patient data^9^ (see Table 1). One way to improve our *in vivo* estimate would be to use new *P. falciparum* data with gametocytaemia measurements over a longer period of time to capture the natural decay of circulating gametocytes, which was not observed in the current VIS data (see Figure 3).

We also predicted the effects of altered gametocyte kinetic parameters on the transmissibility from human to mosquito, focusing on two scenarios: the infectiousness of newly hospitalised malaria patients (i.e. the gametocytaemia when asexual parasitaemia first reaches a hospitalised level of 10^8^ parasites/mL in the model); and the non-infectious period of malaria patients (i.e. the time from the inoculation of infected red blood cells to the time when the gametocytaemia reach a transmission threshold of 10^3^ parasites/mL in the model). We explored how the sexual commitment rate and gametocyte sequestration time influenced the gametocyte level and the non-infectious period. We would like to emphasize that human-to-mosquito transmissibility is determined by both the gametocytaemia level in humans and the relationship between gametocytaemia and the probability of transmission per bite. A reliable prediction of the former is essential but not a sole determinant of transmissibility. Therefore, it is also important to improved our quantitative understanding of the latter, which could be complicated and also determined by the densities of female and male gametocytes^5,6,20^.

Our study has some limitations. The gametocyte population dynamics model, that has been shown to have sufficient complexity to reproduce the clinical observations, is still a rather coarse simplification of the actual biological processes. For example, the model does not assume an adaptive sexual commitment rate^21^, nor does it consider the mechanisms of sexual commitment^22^. Furthermore, the model assumes a constant gametocyte death rate but does not consider other non-constant formulations as have been previously proposed^8^. Another limitation is that we assumed a fixed duration for the asexual life cycle of 42 hours, while previous work by our group suggests that the life cycle may be altered by up to a few hours in response to antimalarial drugs (e.g. artemisinin^23,24^), although it is not known if piperaquine (which was administered in this VIS) has a similar effect.

In conclusion, we have developed a novel mathematical model of gametocyte population dynamics, and demonstrated that it reliably predicts time series data of gametocytaemia. The model provides a powerful predictive tool for informing the design of future volunteer infection studies which aim to test transmission-blocking interventions. Furthermore, the within human host transmission model can be incorporated into epidemiological-scale models to refine predictions of the impacts of various antimalarial treatments and transmission interventions.

## Materials and Methods

### Study population and measurements

The data used in this modelling study are from a previously published VIS^7^ where 17 malaria-naïve volunteers were inoculated with *P. falciparum*-infected red blood cells (3D7 strain). The study was approved by the QIMR Berghofer Human Research Ethics Committee and registered with ClinicalTrials.gov (NCT02431637 and NCT02431650). The volunteers were treated with 480 mg piperaquine phosphate (PQP) on day 7 or 8 post-inoculation to attenuate asexual parasite growth and a second dose of 960 mg PQP was given to any volunteer for treatment of recrudescent asexual parasitaemia. All participants received a course of artemether/lumefantrine and, if required, a single dose of primaquine (45 mg) to clear all parasites at the end of the study. Parasitaemia in the volunteers were monitored at least daily since the inoculation.

The data contain the measurements of total parasitaemia (total circulating asexual parasites and gametocytes per mL blood), asexual parasitaemia (circulating asexual parasites per mL blood), gametocytaemia (circulating female and male gametocytes per mL blood) and plasma concentration of PQP, at multiple time points after inoculation (see Figure 1—3 and Figure S2 in the Supplementary Information). Further details about the VIS are given in^7^. We emphasize that the data used in model fitting is the total parasitaemia (from the first measurement to the time before any treatment other than PQP) and the other data (i.e. asexual parasitaemia and gametocytaemia) are used to validate the model.

### Gametocyte dynamics model

The model extends the published models of asexual parasite life cycle^18,25^ by incorporating the development of gametocytes. The model is comprised of three parts describing three populations of parasites: asexual parasites (*P*), sexual parasites (*P*_*G*_) and gametocytes (G). Note that biologically speaking the phrase “sexual parasites” includes gametocytes, but in order to model the sexual parasites which are indistinguishable from asexual parasites under a microscope, we treat those sexual parasites and gametocytes as two different populations. A schematic diagram of the development of those populations based on current knowledge^11,22^ is shown in Figure 5.

**Figure 5:**
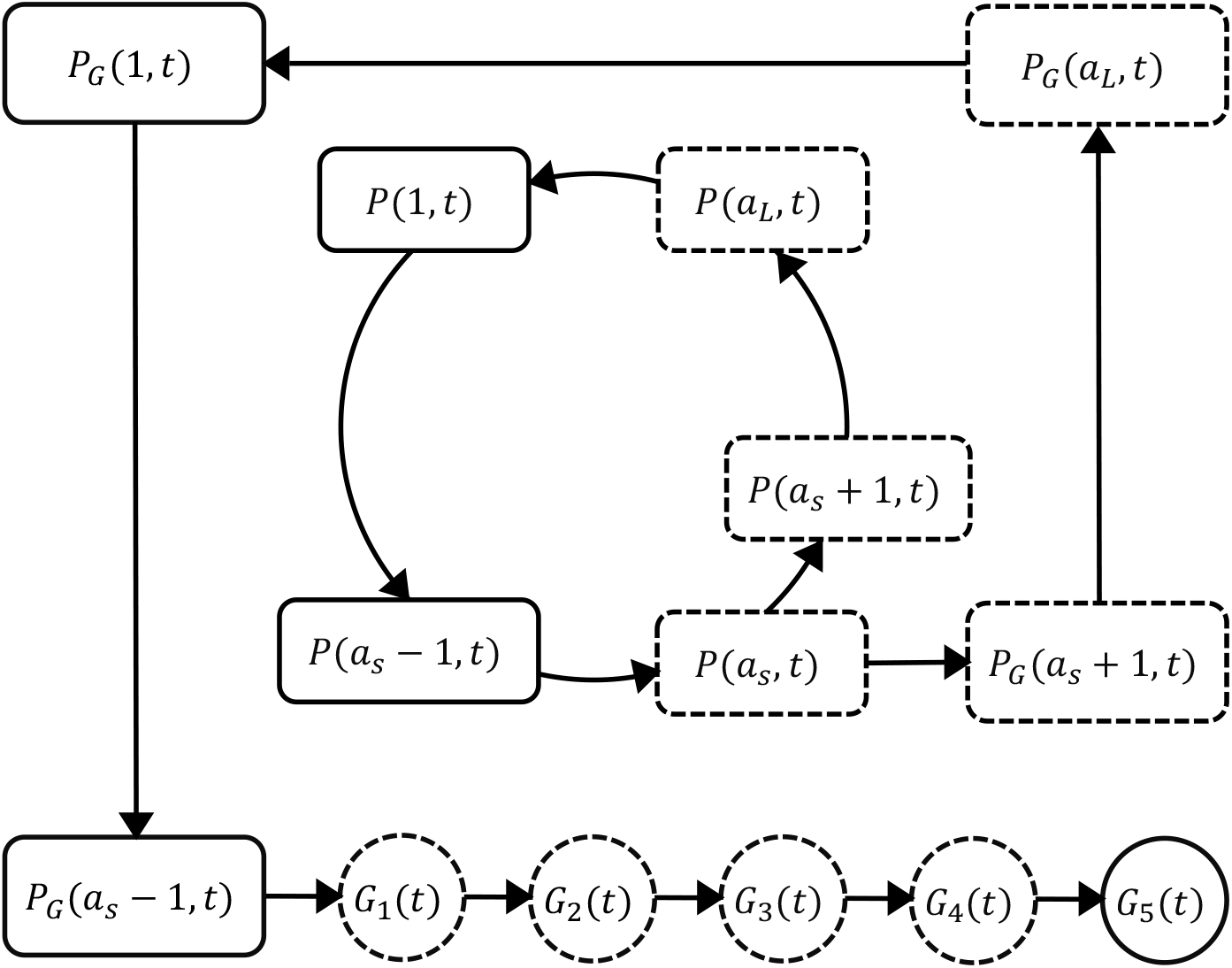
Schematic diagram showing the model compartments and transitions. Square compartments in the inner loop represent the asexual parasite population which follows a life cycle of maturation and replication every *a*_*L*_ hours. Sexual commitment occurs from age *a*_*S*_ and a fraction of asexual parasites become sexual (the bigger square compartments in the outer loop) and eventually enter the development of stage I—V gametocytes (the round compartments). The compartments with a dashed boundary are sequestered to tissues and thus not measurable in a blood smear. The notation of each compartment is consistent with those in model equations and is explained in the main text.

Asexual parasites develop and replicate in the red blood cells (RBCs) till cell rupture at the end of each life cycle and the released free parasites (merozoites) can initiate new life cycles if successfully invading susceptible RBCs. At the time of inoculation (i.e. *t* = 0 hours in the model), we define the inoculum size to be *P*_*init*_ and assume the age distribution of inoculated parasites is Gaussian with mean *μ* and standard deviation *σ*. As time increments by one hour, the asexual parasites of age *a* at time *t* (denoted as *P*(*a*, *t*)) follow the iterative equation:

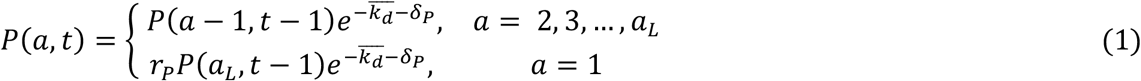

where 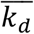 represents the average rate of asexual parasite killing by PQP and *δ*_*P*_ is the rate of asexual parasite death due to processes other than PQP. 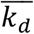 is approximated by the average of *K*_*d*_(*t* − 1) and *K*_*d*_(*t*) and *K*_*d*_(*t*) = *K*_*max*_*C*(*t*)^γ^/(*C*(*t*)^γ^ + *EC*_50_^γ^) where *K*_*max*_ is the maximum killing rate and *EC*_50_ is the PQP concentration at which half maximum killing is achieved. *C*(*t*) is the PQP concentration which is simulated by a pharmacokinetic model introduced below. *a*_*L*_ is the length of each life cycle and *r*_*p*_ is the parasite replication rate indicating the average number of newly infected RBCs attributable to the rupture of a single infected RBC at the end of its life cycle. Sexual commitment is assumed to occur at the first age of the trophozoite stage (denoted to be *a*_*S*_) and a percentage (*f*) of asexual parasites leave asexual life cycle and start sexual development in the next hour, which is modelled by

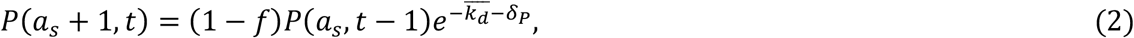

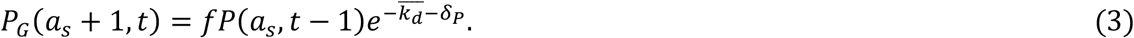

The first equation describes the proportion of parasites remaining in the asexual life cycle while the second equation describes the proportion of parasites becoming sexual parasites (*P*_*G*_). According to Figure 5, the sexual parasites continue the rest of the life cycle and a part of the next life cycle (note that they appear indistinguishable from asexual parasites under microscopy) before becoming stage I gametocytes. The process is modelled by

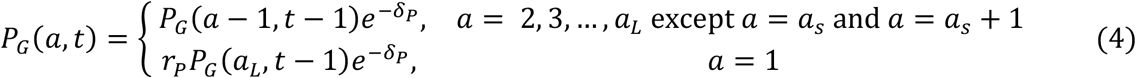

Note that no 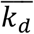 for *P*_*G*_ because PQP does not kill sexual parasites and gametocytes. The changes of the sequestered stage I—IV gametocytes (*G*_1_—*G*_4_) are governed by difference equations

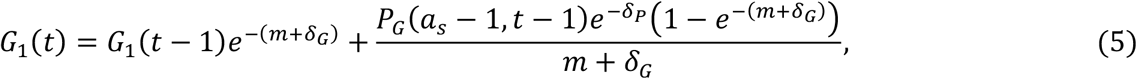

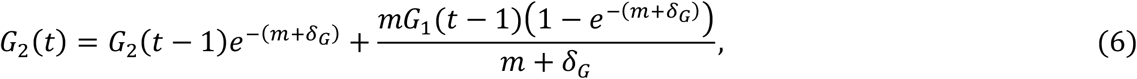

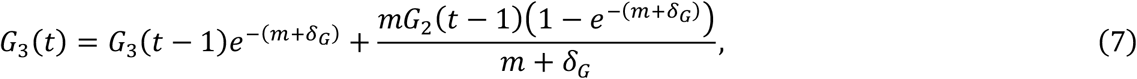

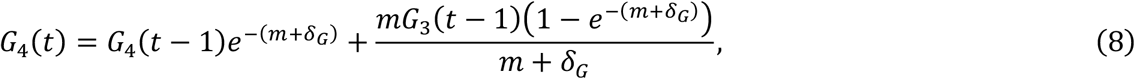

where m is the rate of gametocyte maturation and *δ*_*G*_ is the death rate of sequestered gametocytes. Stage V gametocytes are circulating in bloodstream (and therefore can be measured from the peripheral blood film) modelled by

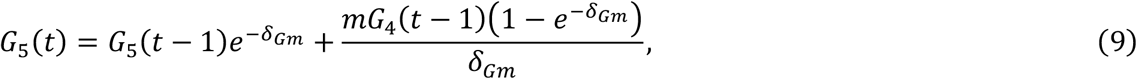

where *δ*_*Gm*_ is the death rate of mature circulating gametocytes.

The total parasitaemia in the model is given by 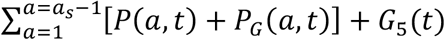, which was fitted to the VIS data. After model fitting, we simulated the asexual parasitaemia 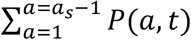 and gametocytaemia *G*_5_(*t*) and compared them with associated data for model validation. Table 2 presents all the model parameters and their units and descriptions.

**Table 2:** Details of the gametocyte dynamics model parameters. The parameter ranges of the prior distributions are chosen based on previous estimates^25^ and biological plausibility. We assumed parasites younger than 25h are circulating and thus fix *a*_*S*_ to be 25h. For 3D7 strain, the life cycle is approximately 39—45h (based on *in vitro* estimates^26^ and *in vivo* estimates by Wockner et al.^17^) and we fix *a*_*L*_ to be 42h. The boundaries of the uniform prior distribution (U) also indicate the lower and upper bounds of the corresponding parameters.

| Parameter | Unit | Description | Prior distribution |
| --- | --- | --- | --- |
| $P_{init}$ | parasites/mL | inoculation size | U(0, 10) |
| $\mu$ | h | mean of the initial parasite age distribution | U(0, 35) |
| $\sigma$ | h | SD of the initial parasite age distribution | U(0, 20) |
| $r_P$ | (unitless) | parasite replication rate | U(0, 100) |
| $k_{max}$ | $h^{-1}$ | maximum rate of parasite killing by PQP | U(0, 1) |
| $EC_{50}$ | ng/mL | half-maximum effective PQP concentration | U(1, 100) |
| $\gamma$ | (unitless) | Hill coefficient for PQP | U(0, 20) |
| $f$ | (unitless) | sexual commitment rate (in percentage) | U(0, 1) |
| $\delta_P$ | $h^{-1}$ | death rate of asexual and sexual parasites | U(0, 0.2) |
| $m$ | $h^{-1}$ | maturation rate of gametocytes | U(0, 0.1) |
| $\delta_G$ | $h^{-1}$ | death rate of sequestered gametocytes | U(0, 0.1) |
| $\delta_{Gm}$ | $h^{-1}$ | death rate of circulating gametocytes | U(0, 0.1) |
| $a_s$ | h | sequestration age of asexual parasites | fixed to be 25 |
| $a_L$ | h | length of life cycle of asexual parasites | fixed to be 42 |

SD: standard deviation; h: hour.

### Pharmacokinetic model of piperaquine (PQP)

In the within-host model, the killing rate *k*_*d*_(*t*) is determined by PQP concentration *C*(*t*) which was simulated from a pharmacokinetic (PK) model introduced in this section. The PK model, provided by Thanaporn Wattanakul and Joel Tarning (Mahidol-Oxford Tropical Medicine Research Unit, Bangkok), is a three-compartment disposition model with two transit compartments for absorption (see schematic diagram in Figure S3, Supplementary Information). Based on Figure S3, the model is formulated to be a system of ordinary differential equations:

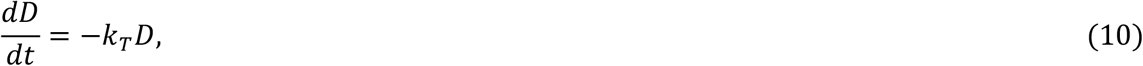

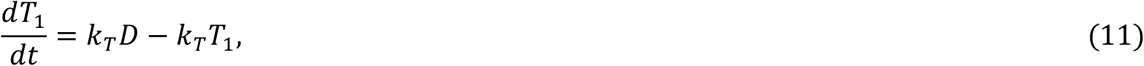

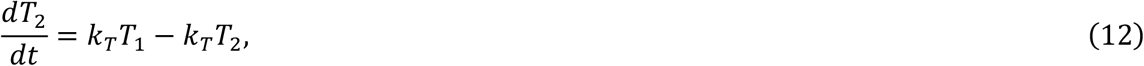

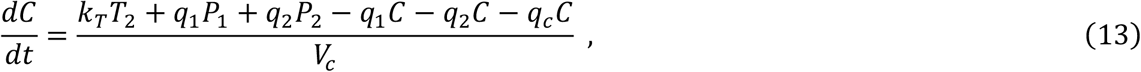

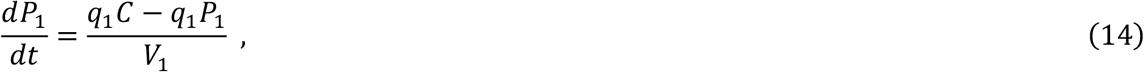

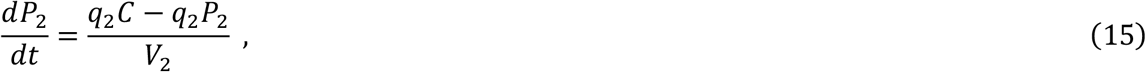

The model parameters were obtained by fitting the model output *C*(*t*) to the PQP concentration data from the VIS in MATLAB (version 2016b; The MathWorks, Natick, MA) using *lsqcurvefit*. The MATLAB code (with detailed comments) are publicly available at https://doi.org/10.26188/5cde4c26c8201. Note that for volunteers whose number of data points is less than the number of parameters in the PK model (such that optimization fails), the parameter values are given by the estimates from another VIS (provided by Thanaporn Wattanakul and Joel Tarning). The best-fit curve and associated parameter values for all volunteers are provided in Figure S2 and Table S1 in the Supplementary Information.

### Fitting the model to parasitaemia data

We took a Bayesian hierarchical modelling approach^27^ to fit the gametocyte dynamics model to the data from all 17 volunteers. In detail, each volunteer has 12 model parameters (i.e. those in Table 2 except *a*_*S*_ and *a*_*L*_; also called the individual parameters) and lower and upper bounds of the parameters are given in Table 2. If denoting the individual parameters to be *θ*_*ind*_ and lower and upper bounds to be *b*_*L*_ and *b*_*U*_ respectively, the following transformations are used to convert the bounded individual parameters to unbounded ones (denoted by *φ*_*ind*_) in order to in order to improve computational efficiency^28,29^:

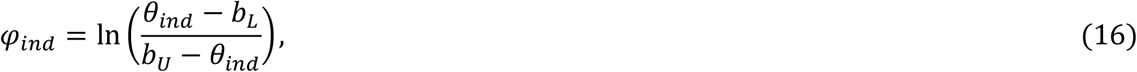

*φ*_*ind*_ obeys a multivariate normal distribution 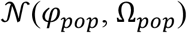 where

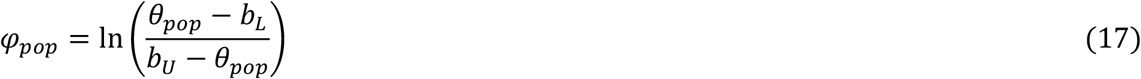

and *θ*_*pop*_ is a vector containing 12 population mean parameters (hyperparameters) corresponding to the 12 gametocyte dynamics model parameters. Ω_*pop*_ is the covariance matrix. For more efficient sampling process, 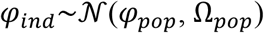 was reparameterised to a non-centred form *φ*_*ind*_ = *φ*_*pop*_ + *ω*_*pop*_*Lη*, where *ω*_*pop*_ is the diagonal standard deviation (SD) matrix whose diagonal elements are the 12 population SD parameters (hyperparameters); *L* is the lower Cholesky factor of the correlation matrix; *η* obeys standard multivariate normal distribution. Note that Ω_*pop*_ = *ω*_*pop*_*LL*^*T*^*ω*_*pop*_ where *LL*^*T*^ is the correlation matrix. The prior distributions for the 12 population mean parameters *θ*_*pop*_ are given by uniform distributions with bounds given in Table 2. The prior distribution for the 12 population SD parameters is standard half-normal and the prior distribution for the lower Cholesky factor of the correlation matrix *L* is given by Cholesky LKJ correlation distribution with shape parameter of 2^29,30^. The distribution of the observed parasitaemia measurements is assumed to be a lognormal distribution with mean given by the model-simulated values and SD parameter with prior distribution of a half-Cauchy distribution with a location parameter of zero and a scale parameter of 5. The distribution for the observed parasitaemia measurements was used to calculate the likelihood function and the M3 method^31^ was used to penalise the likelihood for data points below the limit of detection for the total parasitaemia (10 parasites/mL^7^).

Model fitting was implemented in R (version 3.2.3)^32^ and Stan (RStan 2.17.3)^29^ using the Hamiltonian Monte Carlo (HMC) optimized by the No-U-Turn Sampler (NUTS) to draw samples from the joint posterior distribution of the parameters including the individual parameters (12 parameters for each volunteers) and population mean parameters (12 hyperparameters). Five chains with different starting points (set by different random seeds) were implemented and 1000 posterior samples retained from each chain after a burn-in of 1000 iterations (in total 5000 samples were drawn from the joint posterior distribution). The marginal posterior and prior distributions of the population mean and SD parameters are shown in Figure S4 and S5 in the Supplementary Information. The marginal posterior distributions of the individual parameters for all 17 volunteers are shown in Figure S6—S17 (using violin plots) in the Supplementary Information. For each volunteer, the 5000 sets of individual parameters are used to simulate the gametocyte population dynamics model and generate 5000 simulated model outputs (e.g. 5000 time series of total parasitaemia, asexual parasitaemia or gametocytaemia). The posterior prediction and 95% prediction interval (PI) are given by the median and quantiles of 2.5% and 97.5% of the 5000 model outputs at each time respectively (see Figure 1—3 for example).

The estimates of some key biological parameters (Table 1) were calculated using the 5000 posterior draws of the 12 population mean parameters, i.e. median and 2.5%- and 97.5%-quantile (95% credible interval). The sexual commitment rate was calculated directly by *f*_*pop*_ (the population mean parameter for *f*). Sequestered and circulating gametocyte lifespan were calculated by 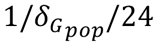 and 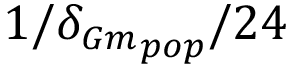 respectively (the factor of 24 converts hours into days). Gametocyte sequestration time was calculated by 4/*m*_*pop*_/24 where 4 indicates four sequestered state (stage I to IV) and 24 converts hours into days. Parasite multiplication factor is calculated by 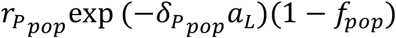 where the term 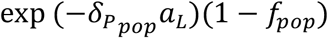 gives the fraction of surviving asexual parasites after death and sexual conversion per life cycle. The gametocyte dynamics model with parameters given by the median estimates of the population mean parameters was used to simulate the two scenarios predicting the dependence of human-to-mosquito transmissibility on the sexual commitment rate and gametocyte sequestration time (Figure 4).

Final analysis and visualization were performed in MATLAB. All computer codes (R, Stan, MATLAB), data and fitting results (CSV and MAT files) and an instruction document (note that reading the document first will make the code much easy to follow) are publicly available at https://doi.org/10.26188/5cde4c26c8201.

## Acknowledgments

We acknowledge useful conversations with David S. Khoury, Deborah Cromer and Miles P. Davenport (Kirby Institute, UNSW Australia, Sydney, Australia) and assistance in drafting the manuscript from Laura Cascales (QIMR Berghofer, Brisbane, Australia). We thank Jörg J. Möhrle, Head of Translation from Medicines for Malaria Venture for his support and permission to use the trial data. The work is supported by an Australian Research Council (ARC) Discovery project (DP170103076) and a National Health and Medical Research Centre (NHMRC) of Australia Project Grant (1025319) and supported in part by the Australian Centre for Research Excellence on Malaria Elimination, funded by the NHMRC (1134989). SGZ is funded by an ARC Discovery Early Career Researcher Award (170100785). JAS is supported by a NHMRC Senior Research Fellowship (1104975). This research was supported by use of the Nectar Research Cloud, a collaborative Australian research platform supported by the National Collaborative Research Infrastructure Strategy (NCRIS).

## Author contributions

P.C., J.A.S., J.S.M. and J.M.M conceived the study; K.A.C. and J.S.M. provided the clinical data; P.C. analyzed the data with contributions from K.A.C., J.A.S., J.S.M. and J.M.M.; P.C. developed the mathematical model; S.Z. developed the statistical method; T.W. and J.T. provided the pharmacokinetic model for piperaquine; P.C. performed model fitting, simulation and analysis; P.C. wrote the first draft of the manuscript; All authors reviewed and commented on the manuscript.

