## Supplementary Information for "Modelling the population dynamics of *Plasmodium falciparum* gametocytes in humans during malaria infection"

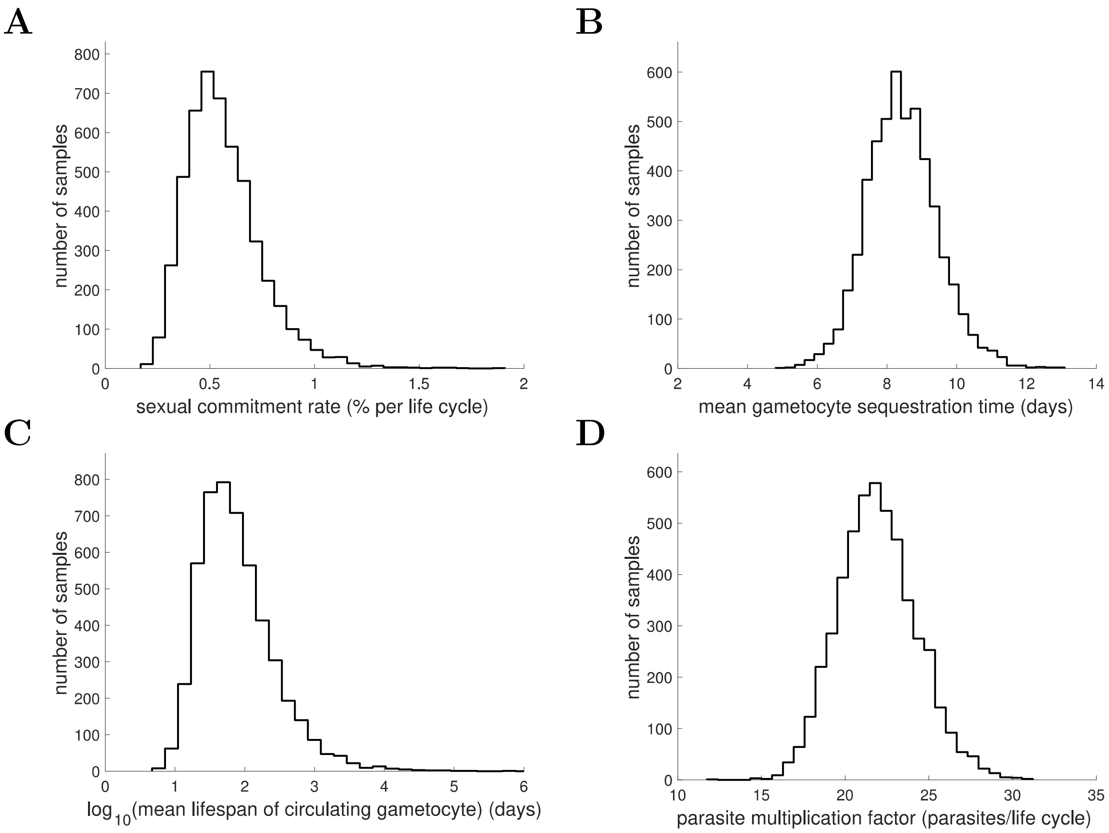


**Figure S1: Marginal posterior distributions of some key biological parameters.** 5000 samples from the posterior parameter distribution are used to generate the figures. Full details about the definitions and expressions of those biological parameters are provided in the Materials and Methods in the main text. Note that a logarithm is taken for the mean lifespan of circulating gametocyte for a better visualisation of the distribution.

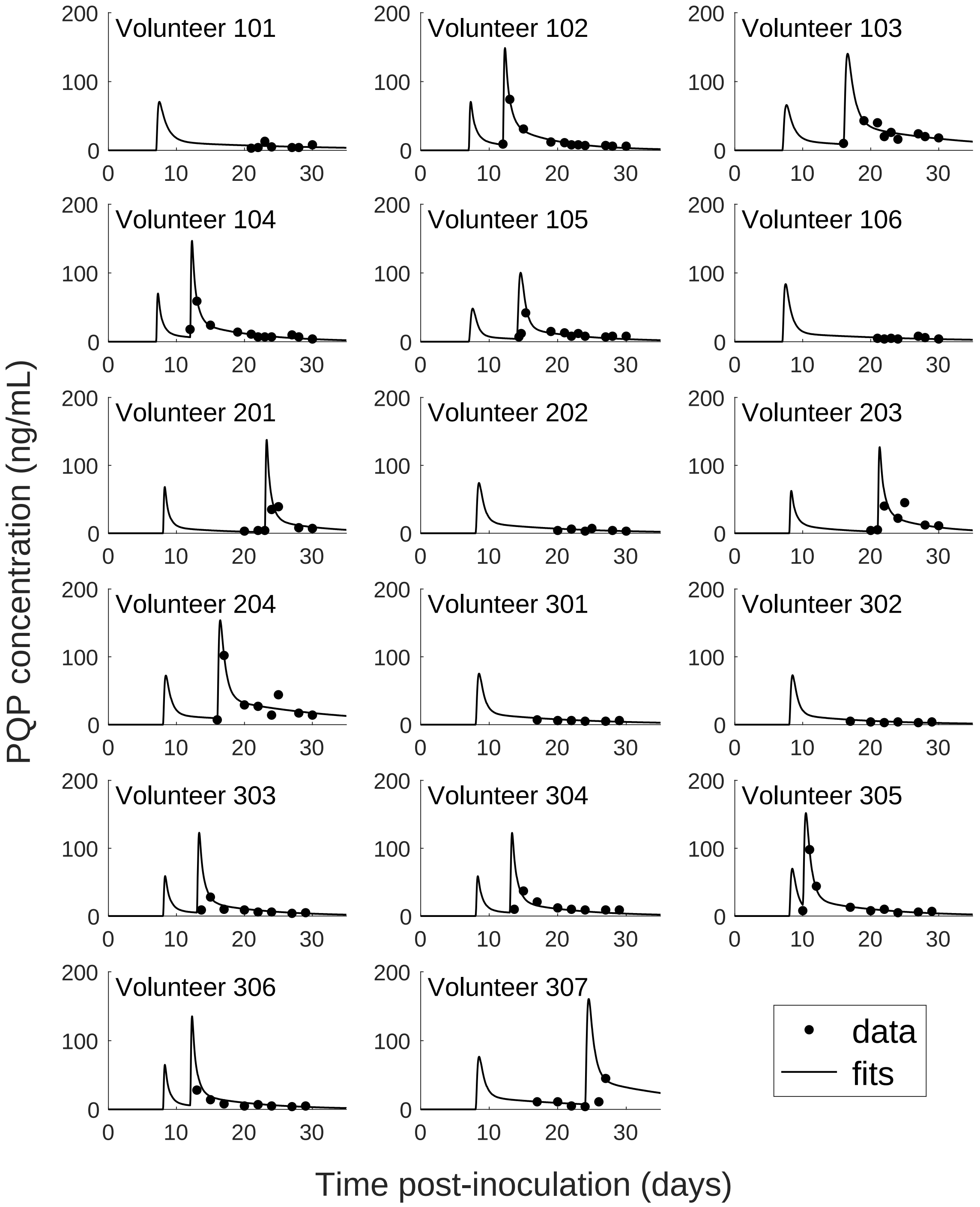


**Figure S2: PK data and model fits of piperaquine (PQP) concentration for all volunteers.** The details of the fitting method are provided in the Materials and Methods in the main text. Best-fit model parameters are given in Table S1.

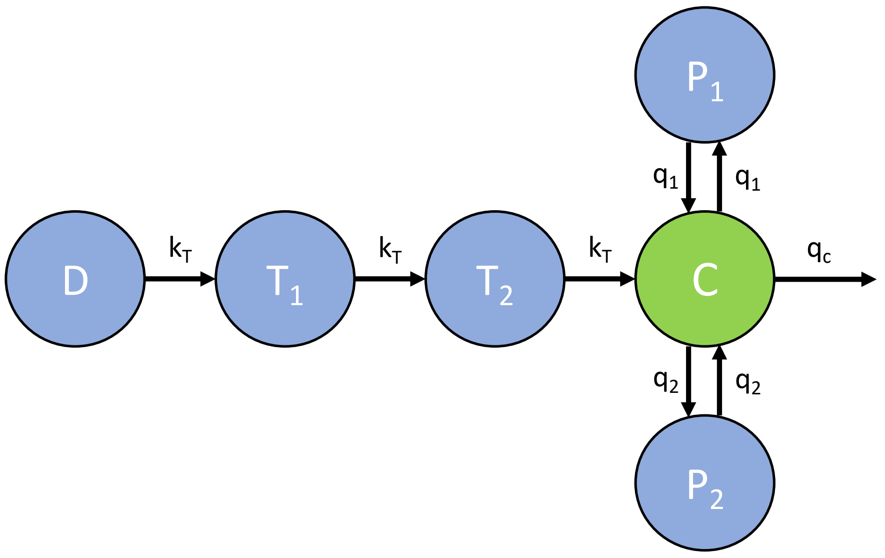


**Figure S3: The pharmacokinetic model of piperaquine (PQP).** The model is a three-compartment disposition model with two transit compartments for absorption. State D represents the dose of PQP. $T_1$ and $T_2$ represent the two transit compartments. C is the central compartment and PQP concentration in this compartment was measured and fitted (which are shown in Figure S2). $P_1$ and $P_2$ represent two peripheral compartments. $k_T$, $q_1$, $q_2$ and $q_c$ are the rates of flow into or out of compartments. Full details and model equations are provided in the Materials and Methods in the main text.

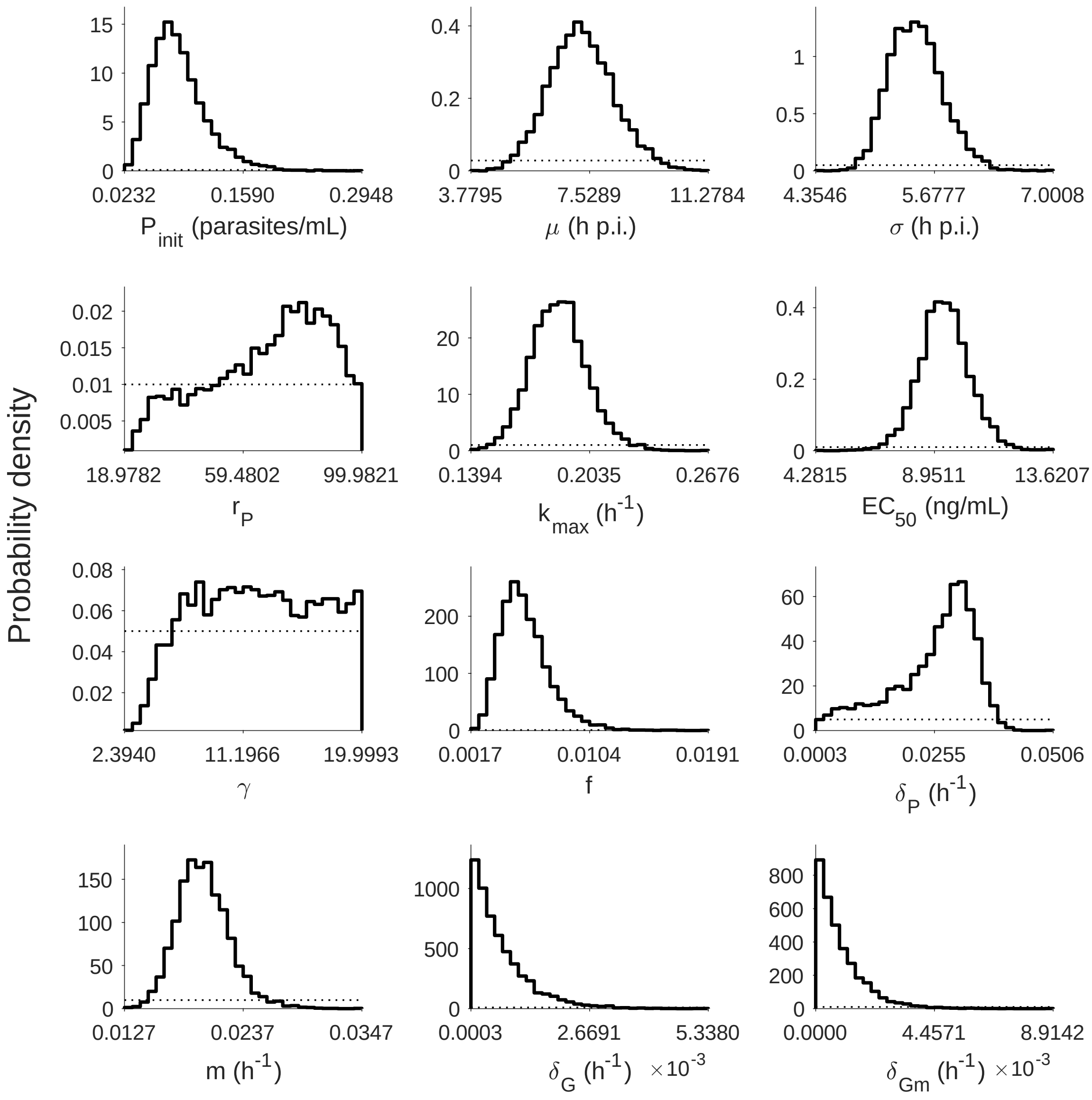


**Figure S4: Marginal posterior distributions for the 12 population mean parameters (hyperparameters).** 5000 samples are used to generate the distributions. The dashed curves indicate the uniform prior distributions. p.i.: post-inoculation. Note that the y-axis is probability density instead of number of samples and the sexual commitment rate f is not converted to percentage. Relevant details of the hyperparameters are provided in the Materials and Methods in the main text.

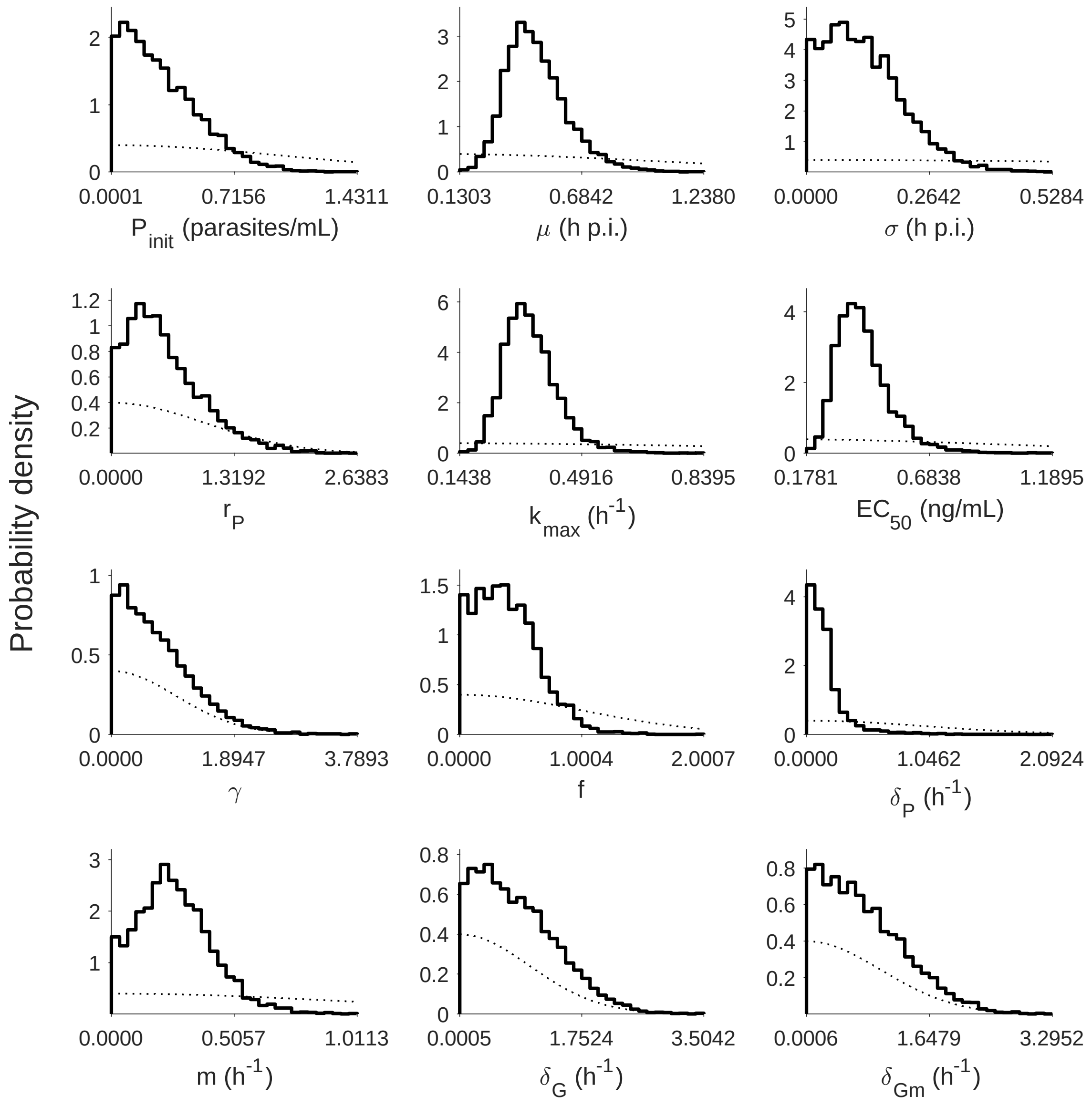


**Figure S5: Marginal posterior distributions for the 12 population SD parameters (hyperparameters).** 5000 samples are used to generate the distributions. The dashed curves indicate the half-normal prior distributions. p.i.: post-inoculation. Note that the y-axis is probability density instead of number of samples and the sexual commitment rate f is not converted to percentage. Relevant details of the hyperparameters are provided in the Materials and Methods in the main text.

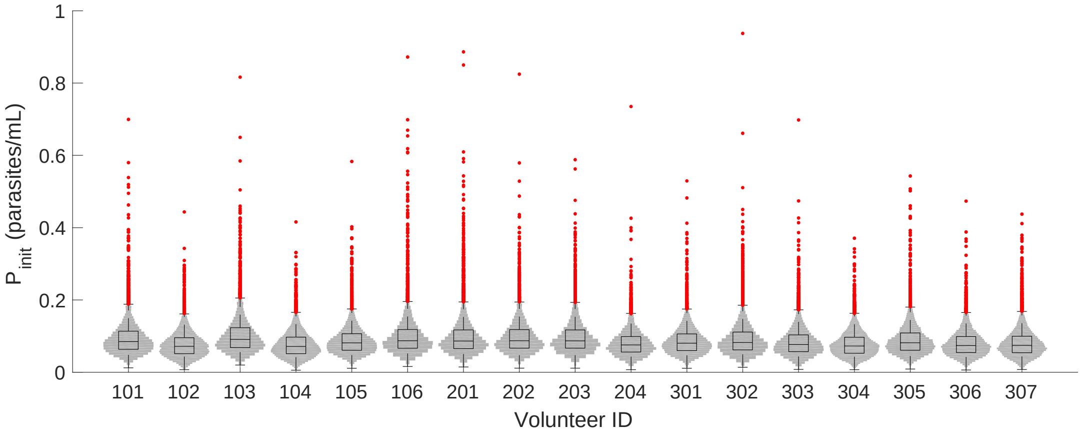


**Figure S6: The marginal posterior distributions of the individual parameter of $P_{init}$ (inoculation size) for all 17 volunteers.** The violin plots (grey area) show the distributions of 5000 posterior samples. Box plots show the 25%, 50% (median) and 75% quantiles with outliers indicated by red dots. Relevant details of the individual parameter are provided in the Materials and Methods in the main text.

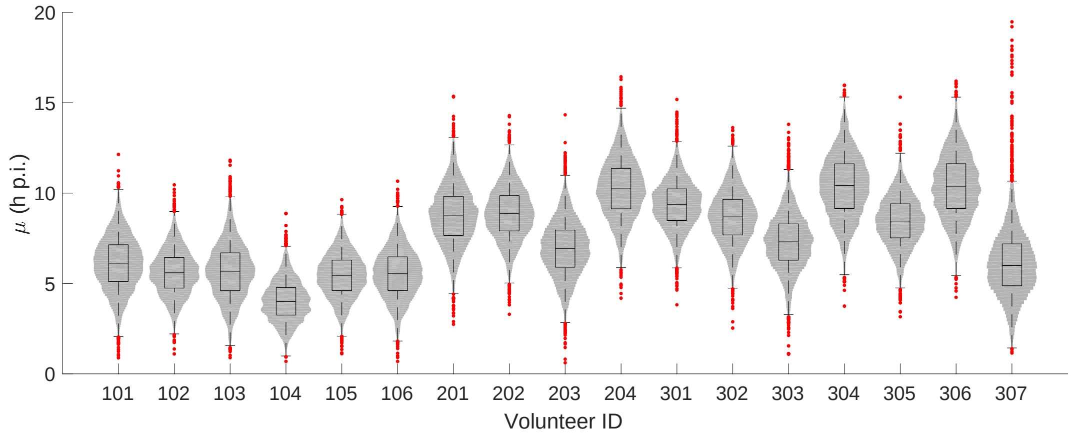


**Figure S7: The marginal posterior distributions of the individual parameter of $\mu$ (mean of the initial parasite age distribution) for all 17 volunteers.** The violin plots (grey area) show the distributions of 5000 posterior samples. Box plots show the 25%, 50% (median) and 75% quantiles with outliers indicated by red dots. Relevant details of the individual parameter are provided in the Materials and Methods in the main text.

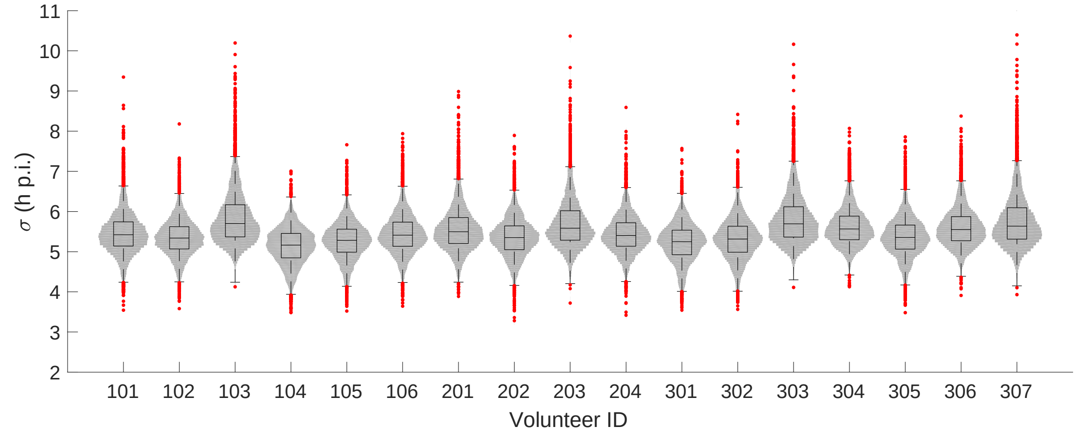


**Figure S8: The marginal posterior distributions of the individual parameter of $\sigma$ (Standard deviation of the initial parasite age distribution) for all 17 volunteers.** The violin plots (grey area) show the distributions of 5000 posterior samples. Box plots show the 25%, 50% (median) and 75% quantiles with outliers indicated by red dots. Relevant details of the individual parameter are provided in the Materials and Methods in the main text.

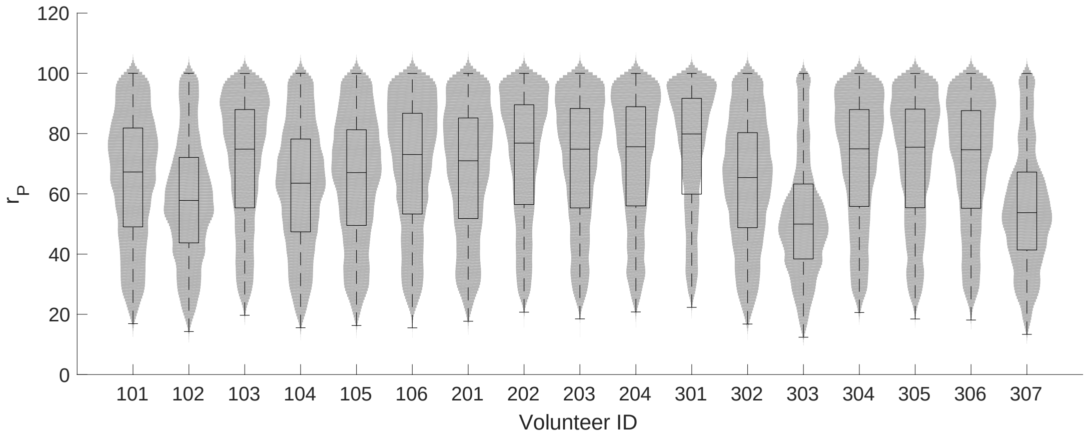


**Figure S9: The marginal posterior distributions of the individual parameter of $r_P$ (parasite replication rate) for all 17 volunteers.** The violin plots (grey area) show the distributions of 5000 posterior samples. Box plots show the 25%, 50% (median) and 75% quantiles with outliers indicated by red dots. Relevant details of the individual parameter are provided in the Materials and Methods in the main text.

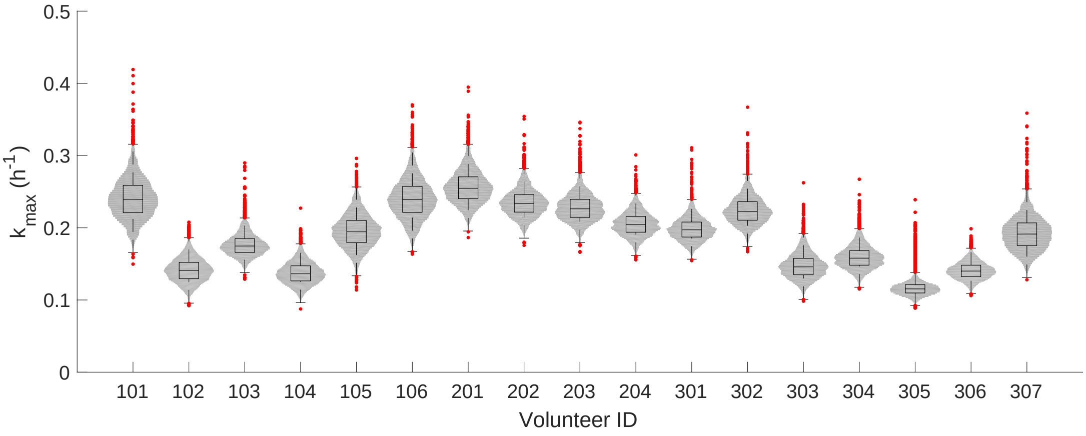


**Figure S10: The marginal posterior distributions of the individual parameter of $k_{max}$ (maximum rate of parasite killing by PQP) for all 17 volunteers.** The violin plots (grey area) show the distributions of 5000 posterior samples. Box plots show the 25%, 50% (median) and 75% quantiles with outliers indicated by red dots. Relevant details of the individual parameter are provided in the Materials and Methods in the main text.

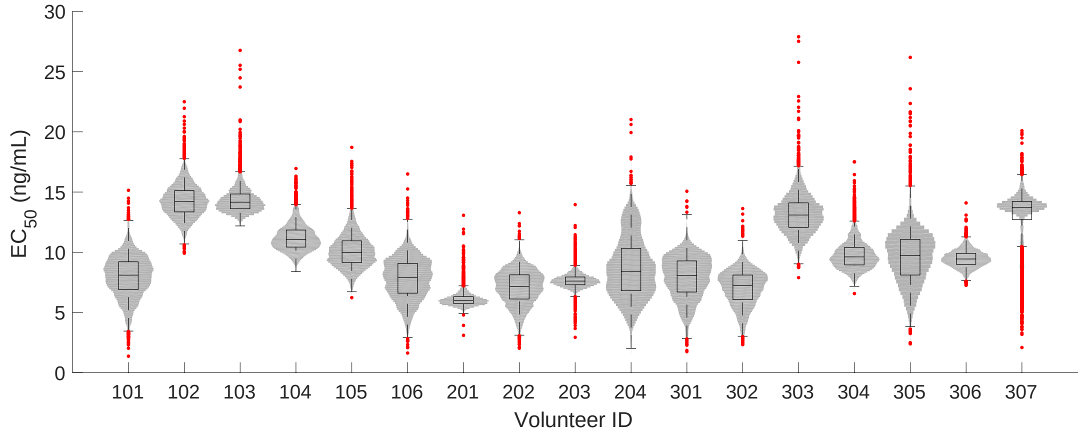


**Figure S11: The marginal posterior distributions of the individual parameter of $EC_{50}$ (half-maximum effective PQP concentration) for all 17 volunteers.** The violin plots (grey area) show the distributions of 5000 posterior samples. Box plots show the 25%, 50% (median) and 75% quantiles with outliers indicated by red dots. Relevant details of the individual parameter are provided in the Materials and Methods in the main text.

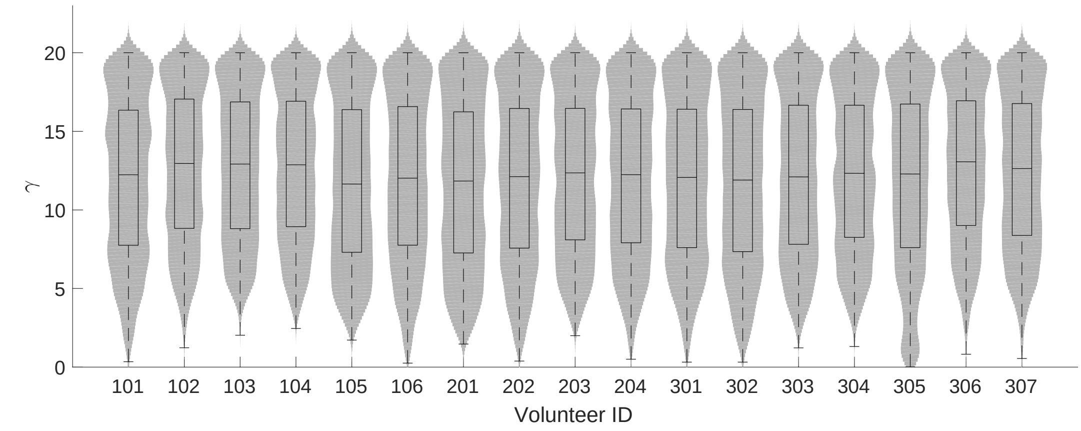


**Figure S12: The marginal posterior distributions of the individual parameter of $\gamma$ (Hill coefficient for PQP) for all 17 volunteers.** The violin plots (grey area) show the distributions of 5000 posterior samples. Box plots show the 25%, 50% (median) and 75% quantiles with outliers indicated by red dots. Relevant details of the individual parameter are provided in the Materials and Methods in the main text.

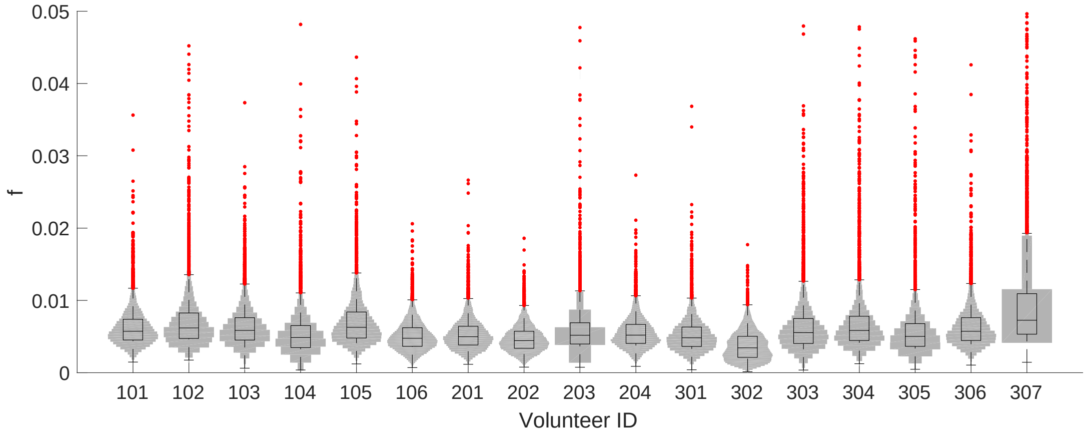


**Figure S13: The marginal posterior distributions of the individual parameter of $f$ (sexual commitment rate; not converted to percentage) for all 17 volunteers.** The violin plots (grey area) show the distributions of 5000 posterior samples. Box plots show the 25%, 50% (median) and 75% quantiles with outliers indicated by red dots. Relevant details of the individual parameter are provided in the Materials and Methods in the main text.

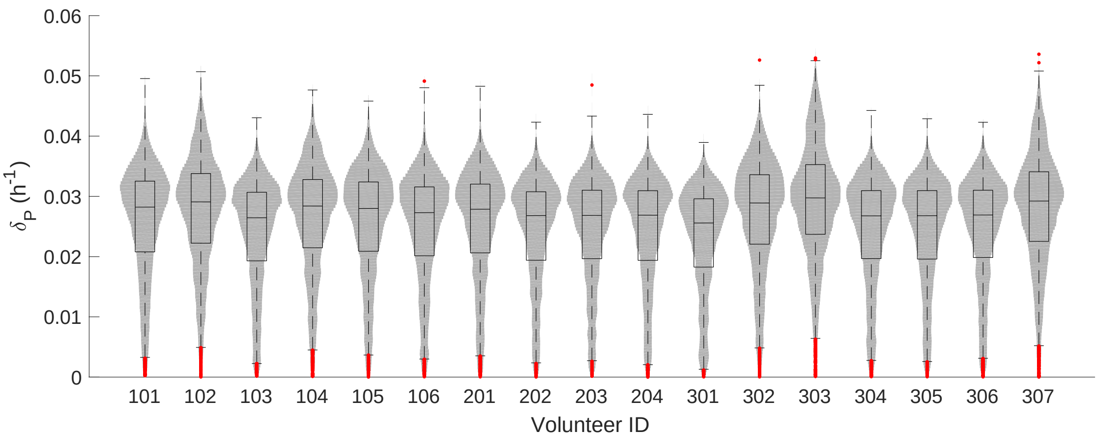


**Figure S14: The marginal posterior distributions of the individual parameter of $\delta_P$ (death rate of asexual and sexual parasites) for all 17 volunteers.** The violin plots (grey area) show the distributions of 5000 posterior samples. Box plots show the 25%, 50% (median) and 75% quantiles with outliers indicated by red dots. Relevant details of the individual parameter are provided in the Materials and Methods in the main text.

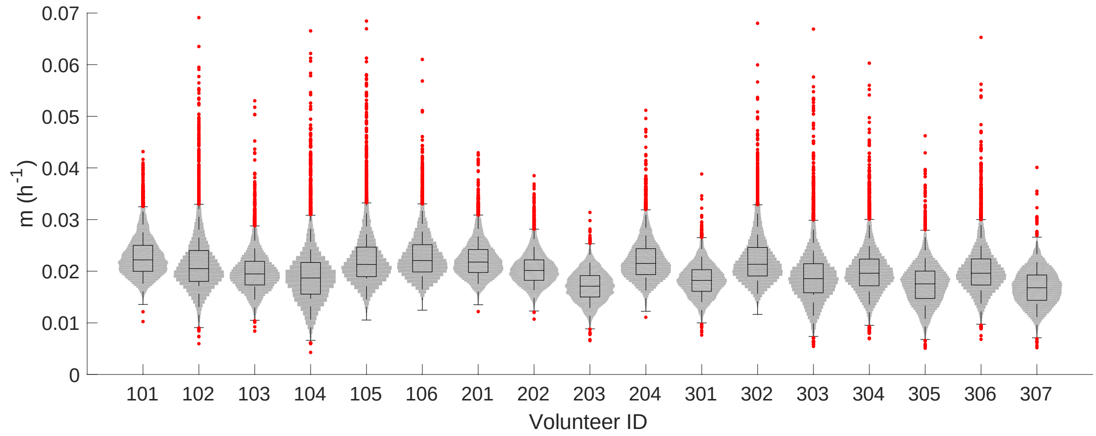


**Figure S15: The marginal posterior distributions of the individual parameter of $m$ (maturation rate of gametocytes) for all 17 volunteers.** The violin plots (grey area) show the distributions of 5000 posterior samples. Box plots show the 25%, 50% (median) and 75% quantiles with outliers indicated by red dots. Relevant details of the individual parameter are provided in the Materials and Methods in the main text.

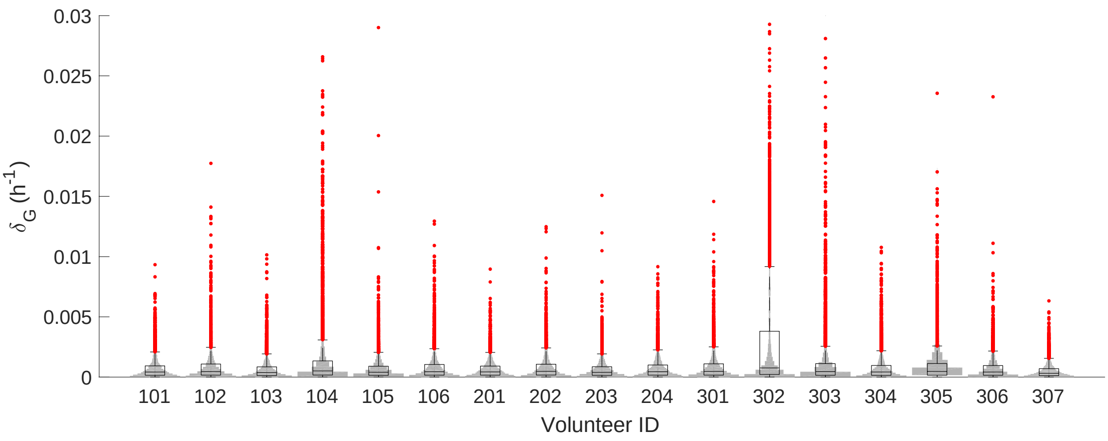


**Figure S16: The marginal posterior distributions of the individual parameter of $\delta_G$ (death rate of sequestered gametocytes) for all 17 volunteers.** The violin plots (grey area) show the distributions of 5000 posterior samples. Box plots show the 25%, 50% (median) and 75% quantiles with outliers indicated by red dots. Relevant details of the individual parameter are provided in the Materials and Methods in the main text.

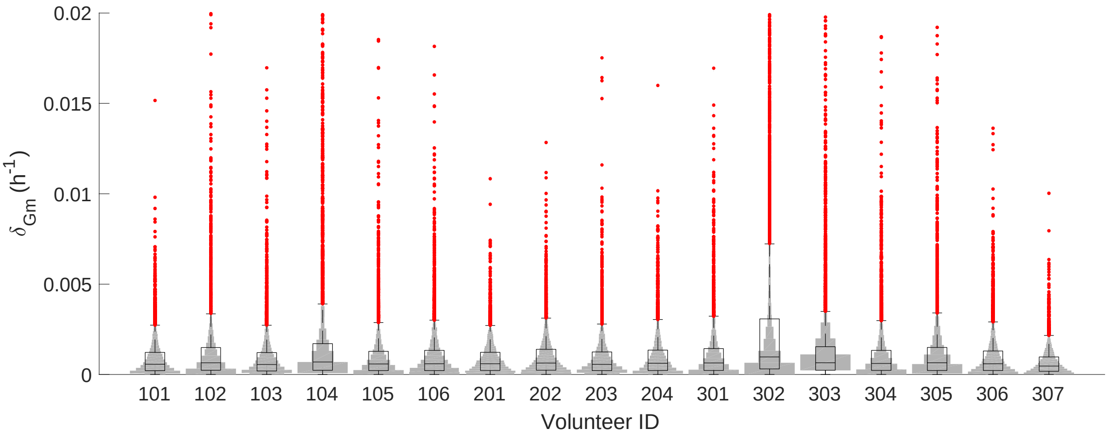


**Figure S17: The marginal posterior distributions of the individual parameter of $\delta_{Gm}$ (death rate of circulating gametocytes) for all 17 volunteers.** The violin plots (grey area) show the distributions of 5000 posterior samples. Box plots show the 25%, 50% (median) and 75% quantiles with outliers indicated by red dots. Relevant details of the individual parameter are provided in the Materials and Methods in the main text.

**Table S1: Best-fit parameter values of the PK model for all 17 volunteers.** Fitting results are shown in Figure S2. The starting values for fitting are $k_T = 1/2.79$, $q_c = 51.4$, $V_c = 804$, $q_1 = 2020$, $V_1 = 3010$, $q_2 = 149$ and $V_2 = 13300$, which are the mean estimates provided by Thanaporn Wattanakul and Joel Tarning (Mahidol-Oxford Tropical Medicine Research Unit, Bangkok). We also assume the parameters are constrained in fitting process by lower bounds of $k_T = 1/3.25$, $q_c = 40.6$, $V_c = 469$, $q_1 = 746$, $V_1 = 2344$, $q_2 = 117$ and $V_2 = 10350$ and upper bounds of $k_T = 1/2.32$, $q_c = 160$, $V_c = 1342$, $q_1 = 5034$, $V_1 = 3986$, $q_2 = 190$ and $V_2 = 17546$. The lower and upper bounds are the 95% quantile of the parameter estimate distributions provided by Thanaporn Wattanakul and Joel Tarning except that the upper bound for $q_c$ is increased from 62.6 to 160 such that the model can reasonably fit to the data. Note that $k_T$ is expressed by the reciprocal of the mean transition time. For Volunteer 202, 301, 302, 307 whose numbers of PK data points are less than the number of parameters in the PK model (such that optimization fails), their "best-fit" parameter values are the starting values given above with some adjustments on $q_c$ (e.g. 71.4 for 202, 61.4 for 301 and 81.4 for 302) which are required to allow the simulated curves to visually capture the data.

| PK parameter (unit) | $k_T$ ($h^{-1}$) | $q_c$ (L/h) | $V_c$ (L) | $q_1$ (L/h) | $V_1$ (L) | $q_2$ (L/h) | $V_2$ (L) |
|---|---|---|---|---|---|---|---|
| **Volunteer 101** | 1/2.43 | 53.7 | 831 | 2240 | 3958 | 118 | 17177 |
| **Volunteer 102** | 1/2.32 | 123.4 | 582 | 849 | 3985 | 163 | 10355 |
| **Volunteer 103** | 1/3.07 | 61.3 | 756 | 2686 | 3910 | 127 | 14346 |
| **Volunteer 104** | 1/2.32 | 134.7 | 488 | 746 | 3986 | 190 | 15007 |
| **Volunteer 105** | 1/3.25 | 160 | 1342 | 5034 | 3986 | 190 | 17546 |
| **Volunteer 106** | 1/2.40 | 60.7 | 667 | 2992 | 3043 | 123 | 16374 |
| **Volunteer 201** | 1/2.32 | 160 | 469 | 746 | 3986 | 190 | 15557 |
| **Volunteer 202** | 1/2.79 | 71.4 | 804 | 2020 | 3010 | 149 | 13300 |
| **Volunteer 203** | 1/2.32 | 141.9 | 1243 | 803 | 3973 | 190 | 11825 |
| **Volunteer 204** | 1/2.32 | 63.3 | 895 | 2169 | 3327 | 165 | 16205 |
| **Volunteer 301** | 1/2.79 | 61.4 | 804 | 2020 | 3010 | 149 | 13300 |
| **Volunteer 302** | 1/2.79 | 81.4 | 804 | 2020 | 3010 | 149 | 13300 |
| **Volunteer 303** | 1/2.45 | 159.4 | 1332 | 779 | 3970 | 190 | 17355 |
| **Volunteer 304** | 1/2.48 | 158.9 | 1339 | 762 | 3981 | 190 | 17290 |
| **Volunteer 305** | 1/2.53 | 120.9 | 1152 | 1905 | 2979 | 123 | 13896 |
| **Volunteer 306** | 1/2.32 | 158.8 | 870 | 761 | 3983 | 190 | 17488 |
| **Volunteer 307** | 1/2.79 | 51.4 | 804 | 2020 | 3010 | 149 | 13300 |
